# Novel Method for Viability Assessment of *Bifidobacterium longum* ATCC 15707 on Non-dairy Foods during drying

**DOI:** 10.1101/403287

**Authors:** Min Min, Susan L. Mason, Grant N. Bennett, Malik A. Hussain, Craig R. Bunt

## Abstract

This study demonstrates a new technique for separating and purifying viable microbes from samples that interfere with viability staining. The viability of *Bifidobacterium longum* ATCC 15707 was assessed using PBDC to separate bacteria from complex non-dairy food matrices and Quantitative Fluorescence Microscopy (QFM) to determine individual cells using LIVE/DEAD *Bac*Light bacterial viability staining. Water agar (3%) was used to retain cells of *B. longum* and offered a lower fluorescence background with *Bac*Light viability staining, compared with fixation on polycarbonate (PC) black membrane. The effect of drying temperatures and non-dairy foods on viability of *B. longum* was assessed. *B. longum* coated on oat, peanut or raisin was separated by filtration, low- and high-speed centrifugation, flotation and sedimentation buoyant density centrifugation. Purified cells were subsequently deposited on water agar for rehydration followed by LIVE/DEAD *Bac*Light viability staining and enumeration. Conventional plate counting was also conducted to compare viability results. Finally, the applicability of this novel method for viability assessment was demonstrated and informative information of cell membrane damages of *B. longum* incorporated onto non-dairy foods during 24 h drying was observed. Viability assessment of *B. longum* coated onto oat, peanut, or raisin was much lower by plate counting compared to viability staining. Drying appeared to have a greater impact when viability was assessed by plate counting compared to viability staining.

**IMPORTANCE:** Enumeration of viable beneficial bacteria from function foods presents a significant bottle neck for product development and quality control. Interference with microscopic and/or fluorescent techniques by ingredients, time required to incubate plated microbes, and the transient nature of the colony forming unit make rapid assessment of viable bacteria difficult. Viability assessment of *Bifidobacterium longum* ATCC 15707 by Percoll Buoyant Density Gradient Centrifugation with LIVE/DEAD *Bac*Light viability staining on water agar (3%) was in agreement with serial dilution enumeration. Without the need for incubation viability assessment by staining provided a more rapid means to assess the impact of drying on the viability of *B. longum* coated onto oat, peanut or raisin.

## INTRODUCTION

Viability of probiotics in food products plays a critical role for delivering health benefits to consumers. Drying is an old method that has been diversely applied to preserve food, disinfect surfaces, prevent pathogen transmission, and to produce and prepare probiotic culture powders (17). During drying, the removal of water can cause desiccation damage to proteins, nucleic acids, lipids and membranes depending on the different levels of drying (20). As a conventional viability determination method, plate counting of colony forming units (CFU) may underestimate numbers of bacteria if bacterial cells are viable but uncultivable, stressed, excessively clumped or inhibited by neighbouring cells or components in the growth media (11). In order to assess viability of probiotic bacteria with a rapid and accurate result, cultivation-independent methods may be better tools to evaluate viability.

Quantitative fluorescence microscopy (QFM) using LIVE/DEAD *Bac*Light Bacterial Viability kit (Invitrogen^®^) in conjunction with epifluorescence microscopy can be used to detect damage of cell membrane for differentiation between “live” (green fluorescent) and “dead” (red fluorescent) cells and enumeration of individual cells. However, the dead cells with intact membranes could be stained with SYTO 9 instead of PI after bacteria were exposed to UVA radiation and EDTA (2). Based on this observation, it would be more suitable to describe SYTO 9 or PI stained cells as green or red fluorescent cells rather than live or dead cells. Besides, some factors such as the bleaching effect of SYTO 9, fluorescence background interference of substrates (22) can also impact results of *Bac*Light viability staining. QFM to determine viability requires low fluorescence background and high signal of green or red fluorescent cells to achieve accuracy and precise of viability assessments (24). Using 0.22 μm polycarbonate (PC) black membranes to trap bacterial cells has been reported by some studies (7, 10, 18, 21). However this filter membrane method could underestimate total cells when environmental samples (5) and non-dairy drink (13) were determined. Because PI fluorescence dye causes background fluorescence and non-specific binding to the matrix of samples (3).

In this study, a new method of cell fixation was developed by trapping cells on water agar (3%). To reduce high fluorescence background and noise that can be caused by food matrix and other debris, Percoll buoyant gradient density centrifugation (PBDC) combined with low and high speed centrifugation was examined in this study. *B. longum* ATCC 15707 was chosen as a model bacterium based on our previous studies that this bacterium exhibited a better survival on tested non-dairy foods during storage at 30 °C and 50% relative humidity (RH) compared to lactobacilli (14).

## RESULTS

### Bouyant density of *B. longum* ATCC 15707 and food fractions

Using a bright field microscopy to exam each buoyant gradient solution that was dried on a glass slide, *B. longum* ATCC 15707 was found in a buoyant gradient layer of 1.052 g mL^-1^, while fractions of oat, peanut and raisin were found in buoyant gradient layers between 1.121 and 1.061 g mL^-1^. Therefore, four Percoll buoyant gradient solutions (1.052, 1.061, 1.075 and 1.121 g ml^-1^) were used for preparing samples using PBDC.

### Bacterial viability after PBDC and stored at 4 °C overnight

An optimal centrifuge time of 60 min giving good separation was determined where *B. longum* ATCC 15707 cells floated to the top layer of the Percoll gradient solutions and few food fractions were observed. However, the PBDC process and image acquisition process times for each batch of samples meant storing at 4 °C and assessment of potential viability loss during storage needed to be determined. A reduction of green fluorescent cells after at 4 °C overnight was found (Fig. 1). Decreased viability from log 8.1 to log 8.0 cells mL^-1^ was observed immediately following the PBDC process, and the viability then further reduce to log 7.9 cells mL^-1^ after storage overnight at 4 °C. This result might indicate that PBDC caused mild damage to cell membranes of *B. longum* ATCC 15707 cells and their viabilities were not significantly decreased. However, these cells while stored at 4 °C overnight led to a significantly declined in viability of log 0.2 cells mL^-1^ (*p* < 0.05) compared with initial viability of log 8.1 cells mL^-1^. Compared with other centrifugation methods that were attempted to purify *B. longum* ATCC 15707 cells from culture coated foods before using PBDC, the PBDC method showed a better separation of cells from food fractions and a lower background fluorescence and noise in QFM, and hence was used in the next experiment of studying the influence of temperatures and food matrix on cell membrane integrity of *B. longum* ATCC 15707 for 24 h drying.

**FIG 1.**
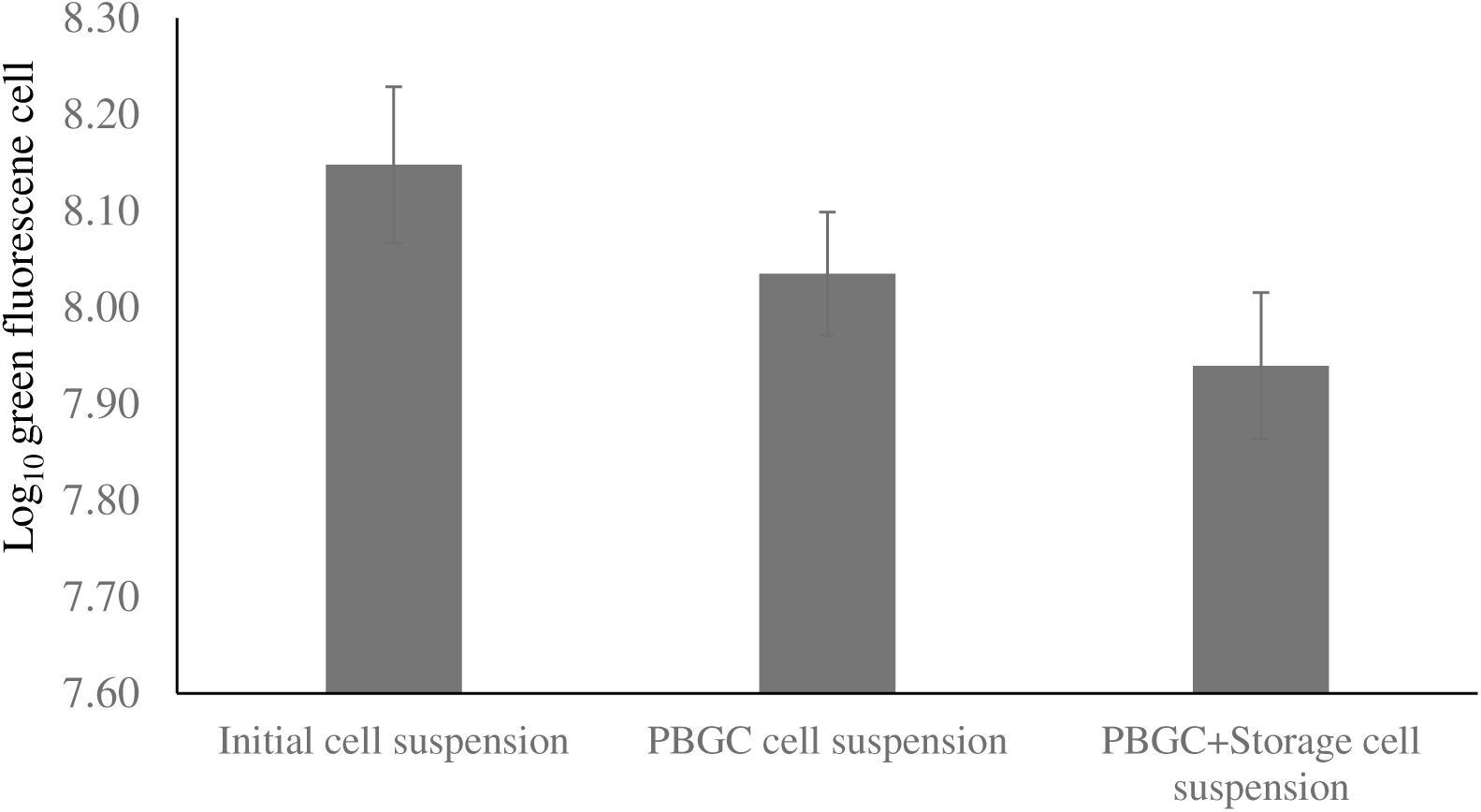
Mean numbers (Log_10_ cells mL^-1^) of green fluorescent cells of *B. longum* ATCC 15707 determined; initially, after Percoll Buoyant Gradient Density Centrifugation (PBDC), and after storage at 4 °C overnight. Error bars are the standard error of the mean (n=5).

### Specificity of QFM

Optimal contact time for staining *B. longum* ATCC 15707 on water agar (3%) was 20 min (in dark at room temperature) so that water in 50 µL of a sample could evaporate or absorb into the agar, thus improving the sharpness of fluorescence images. A minimum cell density of 10^7^ cells mL^-1^ was required for detecting *B. longum* ATCC 15707 cells. An optimum cell density range was between 1 ∼ 3 x 10^8^ cells mL^-1^, representing a range of 50 ∼ 200 green or red fluorescent cells per 10 images (640 x 480 pixels per image).

In this study, water agar (3%) was used for bacterial cell fixation instead of 0.22 µm Cyclopore^TM^ track etched polycarbonate (PC) black membrane (Whatman^®^, USA). When comparing background fluorescence and viability of the cells trapped on the membrane or the agar (Fig. 2), water agar base appeared to have less interference of red fluorescence background than PC black membrane, and hence had a higher ratio of signal-to-noise. That is helpful for automatic green or red fluorescent cell counting via Image J. Additionally, loading and staining cells directly on the agar was easier than filtering, staining and rinsing cells on the membrane. A higher ratio of green or red fluorescent cells exposed to room temperature for 6 h was also observed on the agar than the cells fixed on the membrane. Considering the above advantages of fixing cells on the agar, this method could facilitate preparation and assessment of viability for sample batches and hence could reduce measurement errors between samples.

**FIG 2.**
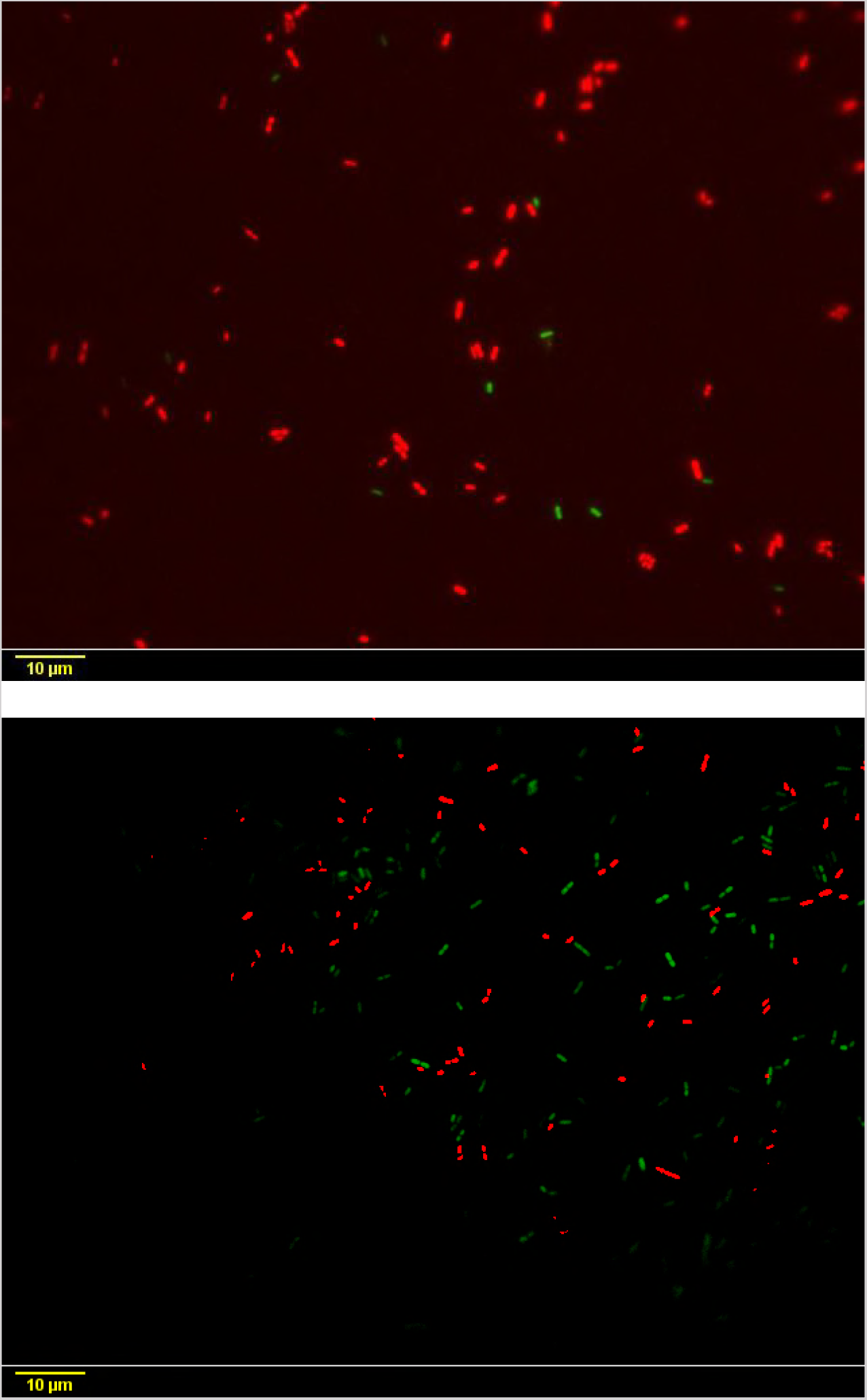
Fluorescence microscopy images of *B. longum* ATCC 15707 cells deposited on a PC black membrane (Top) or water agar (3%) (Bottom) observed with x 1,000 magnification.

### Accuracy of QFM

The calibration curves of automatic cell counting or manual cell counting against percent live cells of *B. longum* ATCC 15707 were plotted in Fig. 3. Green/red fluorescence ratio for live bacteria between 0 % and 100 % was determined manually and compared to the result from automatic cell counting via Image J. These methods generated two calibration curves that showed a linear relationship between % live bacteria and ratio of green/red fluorescent cells (R^2^ = 0.9305 for automatic cell counting; R^2^ = 0.9405 for manual cell counting).

**FIG 3.**
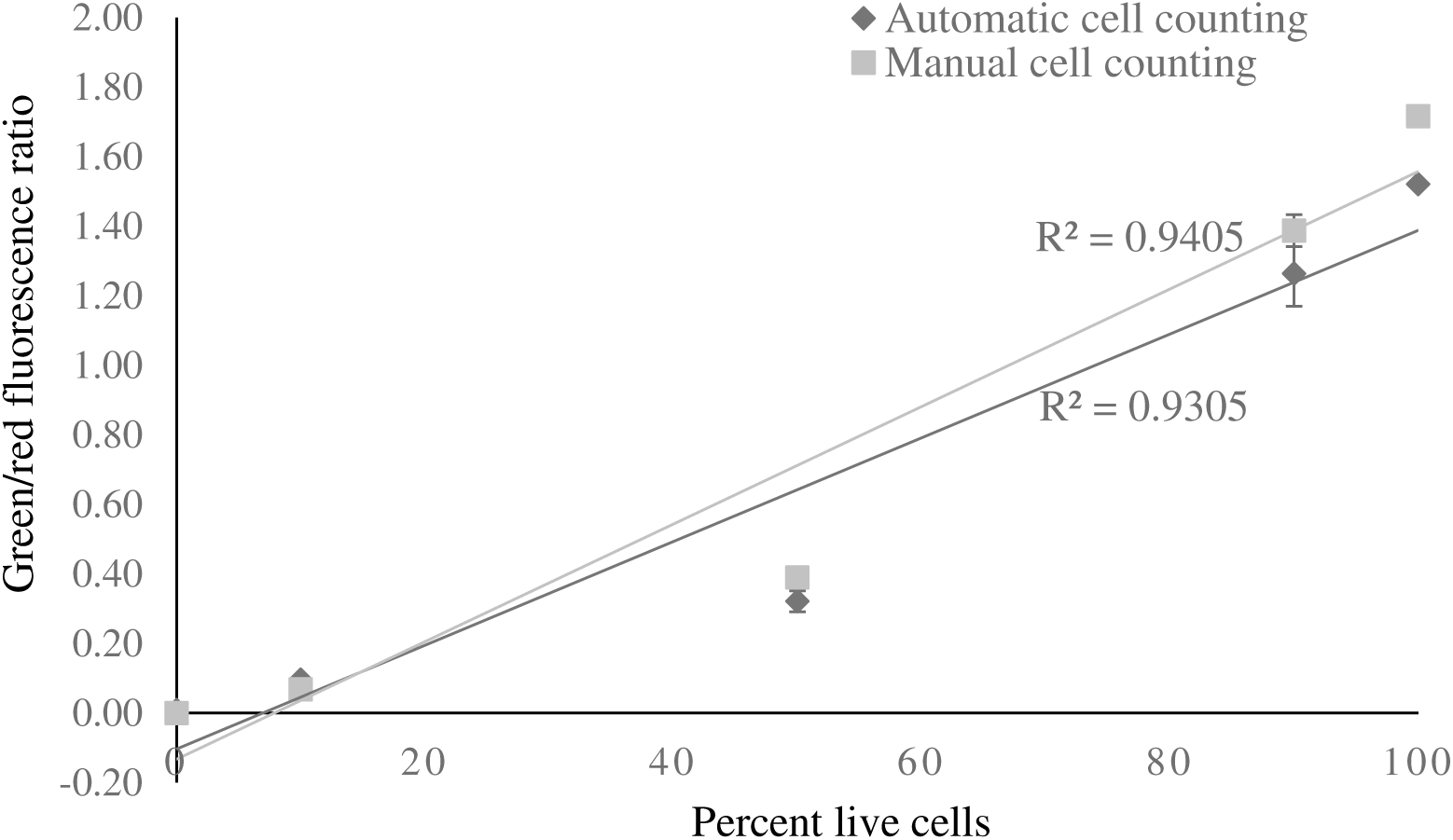
Calibration curve of the relationship between % live bacteria and ratio of green/red fluorescence cells. Error bars are the standard error of the mean (n=3).

### Repeatability of QFM

Six replicates of each *B. longum* ATCC 15707 culture were used on three consecutive days for enumerating green fluorescent cells and cultivable cells by QFM and plate counting, respectively. The repeatability result for intra-day and inter-day variations between the replicates were shown in Table 1. For the intra-day variation, the determined values of coefficient of variance (CV) by QFM method were between 0.792 % and 1.522 %. The CV range of 0.994 % to 1.397 % determined by plate counting was noticed. On the other hand, when considering variability of the two methods across different days, the QFM method showed a much smaller value (% CV=1.152) than plate counting (% CV=5.483). These results may indicate that both methods can produce a relatively low variance of viability determined within a day, but plate counting may have measured with a relatively high variance viability between different days, probably due to variations between viability of different cultures, growth media and incubation conditions.

**Table 1.** Repeatability for determination of viability of *B. longum* ATCC15707 using Quantitative Fluorescence Microscopy (QFM) and plate counting.

| Method | Variation | Mean (Log<sub>10</sub> viability) | SEM(n=6) | % CV |
| --- | --- | --- | --- | --- |
| Quantitative Fluorescence Microscopy | Intra-day | 8.003 | 0.050 | 1.522 |
|  |  | 7.963 | 0.026 | 0.792 |
|  |  | 7.958 | 0.038 | 1.154 |
|  | Inter-day | 7.975 | 0.022 | 1.152 |
| Plate counting | Intra-day | 6.599 | 0.038 | 1.397 |
|  |  | 6.465 | 0.036 | 1.376 |
|  |  | 5.847 | 0.024 | 0.994 |
|  | Inter-day | 6.304 | 0.081 | 5.483 |

### Use of QFM and PBDC for viability assessment of *B. longum* ATCC 15707 coated on non-dairy foods for 24 h drying at two temperatures

In general, viability declined when *B. longum* ATCC 15707 coated on oat, peanut and raisin (Fig. 4.a-c) during 24 h of drying when determined by both QFM and plate counting methods. Compared with the microscopic method, lower average viabilities of log 2.7 CFU g^-1^, log 2.9 CFU g^-1^ and log 3.8 CFU g^-1^ were determined by counting colonies where *B. longum* ATCC 15707 cells were recovered from culture coated oat, peanut and raisin samples, respectively, at 0 and 2 h of drying at 20 °C. When the drying time increased to 24 h (for all temperatures), a statistically significant reduction in CFU g^-1^ was found (*p* < 0.05), but green fluorescent cell numbers appeared to be consistent. It is worth considering that the use of two enumeration techniques can reveal the difference of viability, indicating the cellular status of *B. longum* ATCC 15707 cells possibly changed from viable to viable but not culturable in a mild drying process.

**FIG 4.a.**
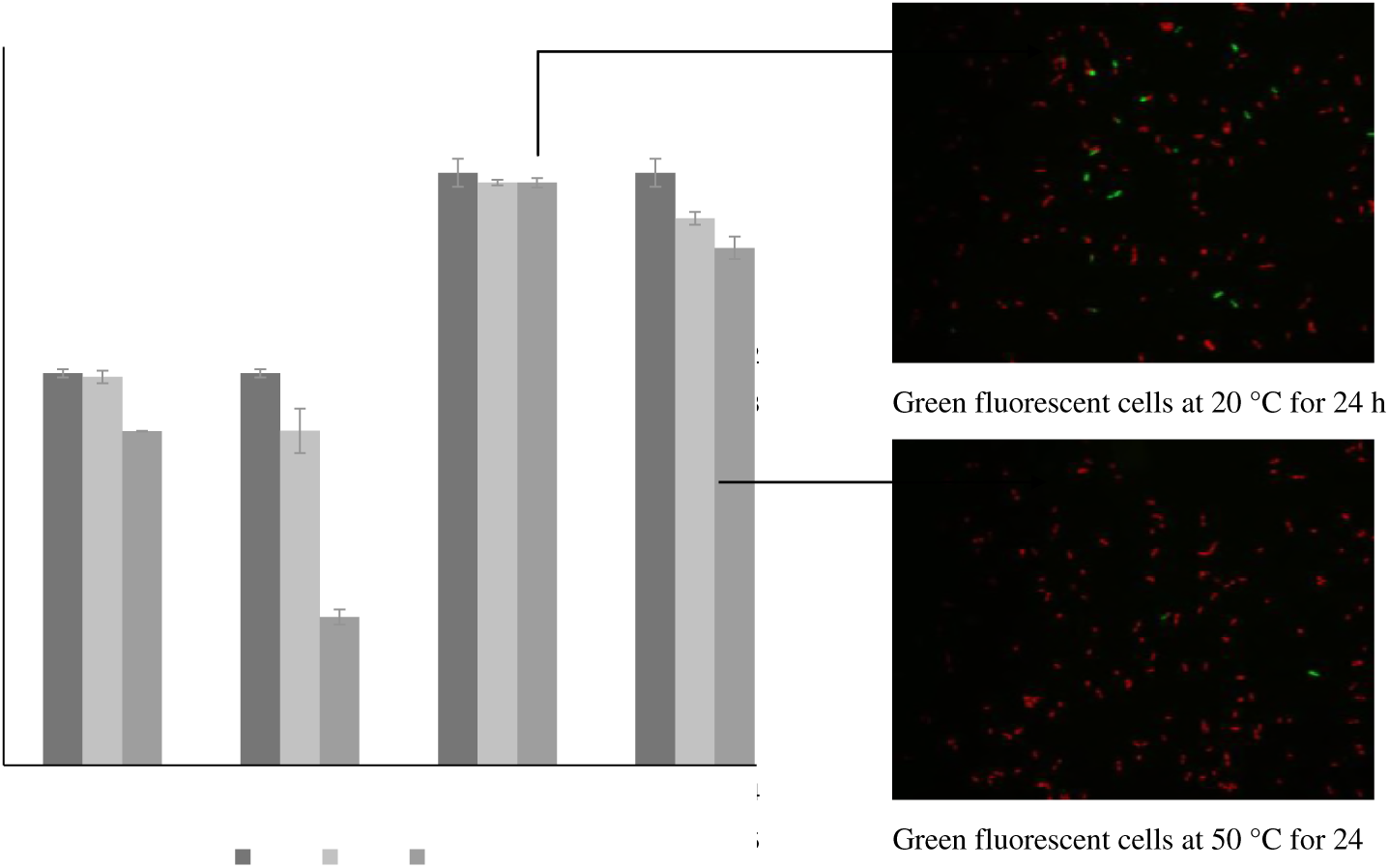
Viability of *B. longum* ATCC 15707 coated on oat and stored at 20 °C and 50 °C for 0, 2 and 24 h. Green and red fluorescent cells were observed in images with x 1,000 magnification. Error bars are the standard error of the mean (n=3).

**FIG 4.b.**
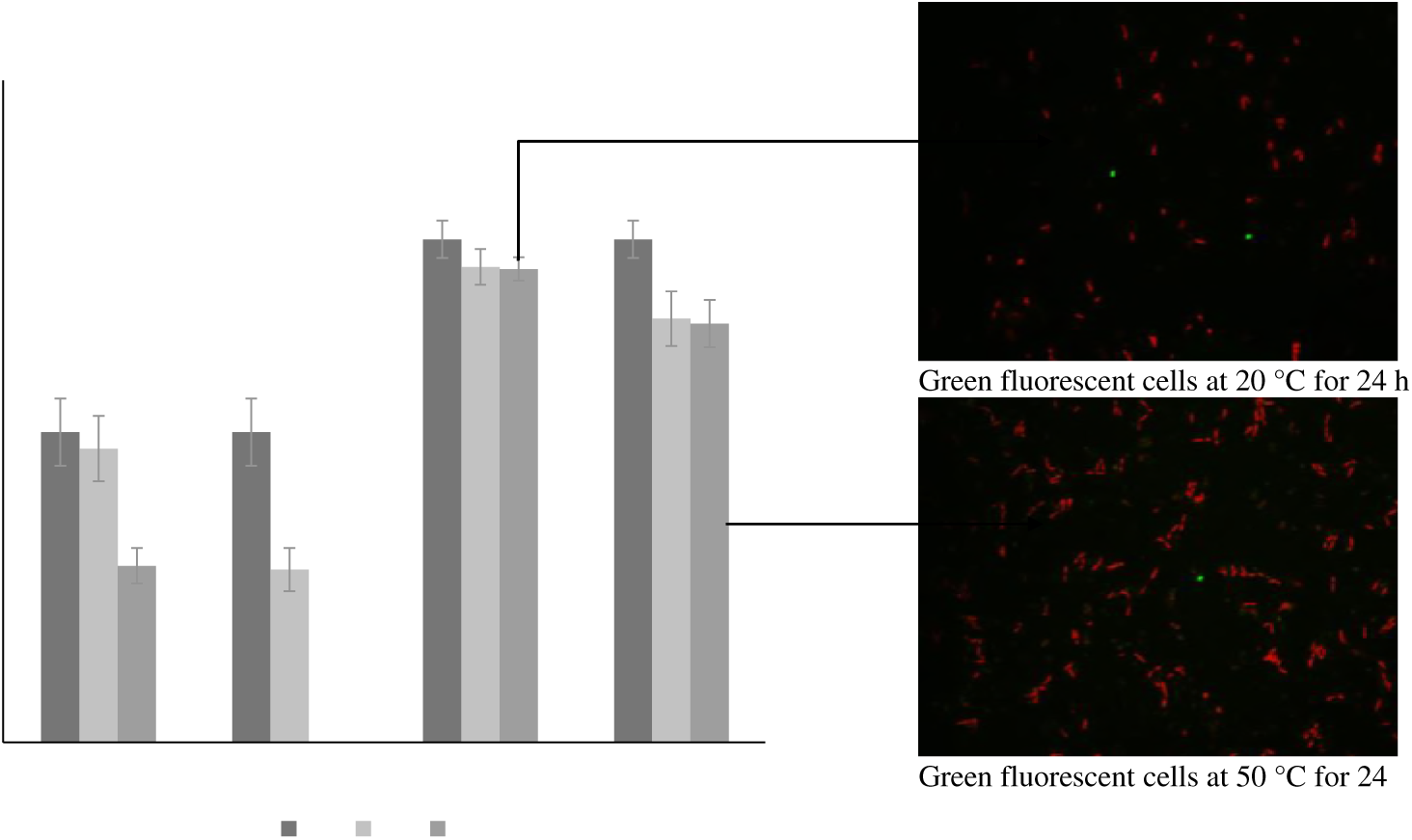
Viability of *B. longum* ATCC 15707 coated on peanut and stored at 20 °C and 50 °C for 0, 2 and 24 h. Green and red fluorescent cells were observed in images with x 1,000 magnification. Error bars are the standard error of the mean (n=3).

**FIG 4.c.**
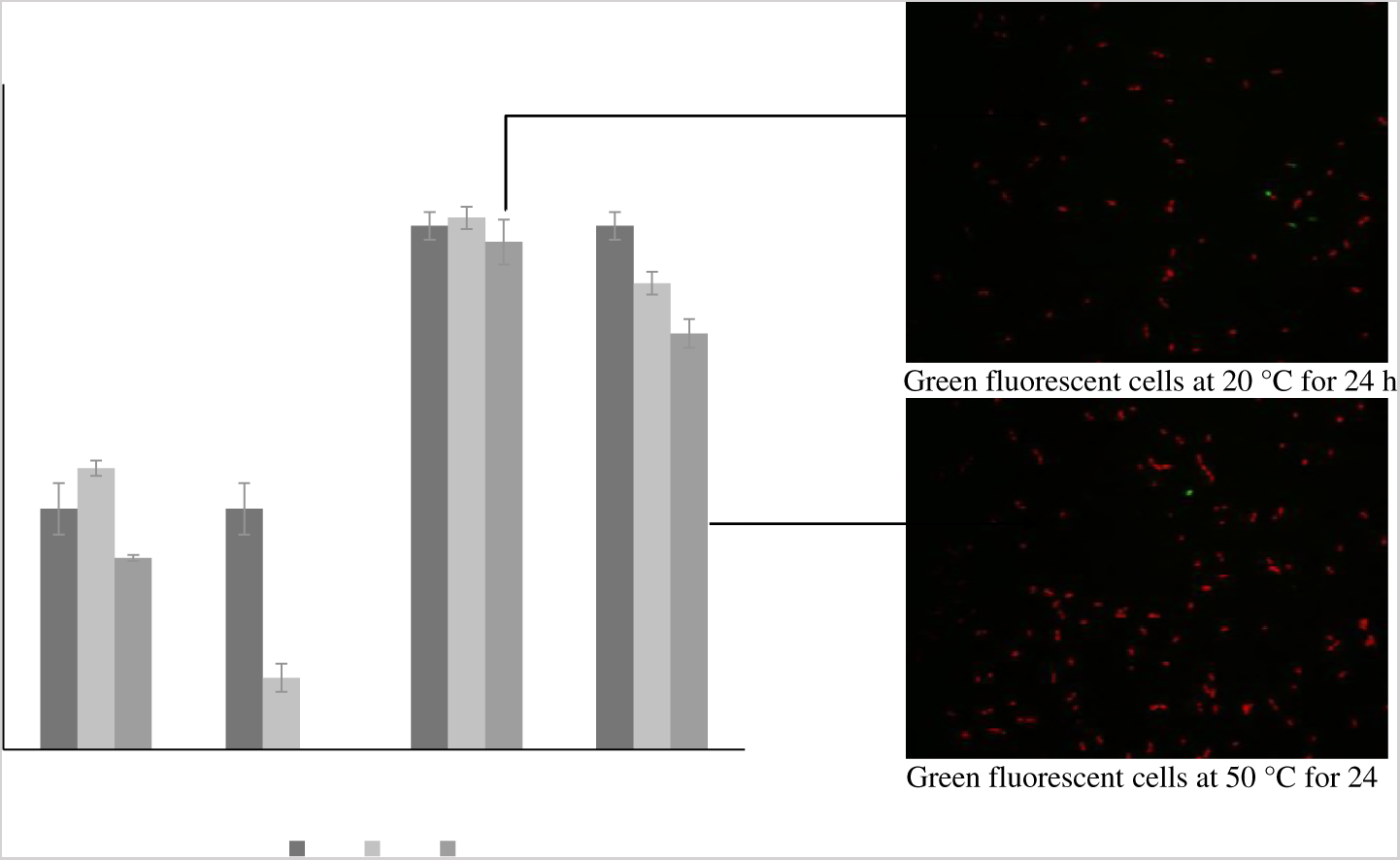
Viability of *B. longum* ATCC 15707 coated on raisin and stored at 20 °C and 50 °C for 0, 2 and 24 h. Green and red fluorescent cells were observed in images with x 1,000 magnification. Error bars are the standard error of the mean (n=3).

When coated samples were dried at 50 °C, a significant decline of viability was detected by both methods (*p* < 0.05). The green fluorescent cells decreased from log 8.2 cells g^-1^ to log 7.0 cells g^-1^ on oat, log 7.6 to log 6.7 cells g^-1^ on peanut, and log 7.8 to log 6.2 cells g^-1^ on raisin after 24 h, while compared with the enumeration result determined by plate counting, the colonies of *B. longum* ATCC 15707 coated on peanut and raisin were not detectable at the end of 24 h, and only log 2.0 CFU g^-1^ was determined on the oat samples. This strongly suggests a different protecting effect of food matrix on cell membrane against a drying process.

## DISCUSSION

Validation of a quantitative fluorescence microscopy (QFM) method for assessing viability bacteria has been applied to *B. longum* ATCC 15707 from a pure culture and subsequently coated on different non-dairy foods after drying at 20 °C and 50 °C. Cell population density is a critical factor affecting green fluorescent cell counts. Low cell density may not provide sufficient numbers of cells for enumeration, and high cell density may induce flocking or overlapping of cells. The minimum detection limit for this QFM was ∼ 10^7^ cells mL^-1^ for *B. longum* ATCC 15707 and an ideal cell density between 1 ∼ 3 x 10^8^ cells mL^-1^ gave 50 to 200 cells per image, which is in accordance with the recommended cell density of 10^8^ cells mL^-1^ for *Bifidobacterium* UCC 35612 (1). A similar study was conducted by measuring viability of *L. plantarum* WCFS1 in 20 images with 200 to 500 cells per image (2,080 by 1,544 pixels) (19). Another study suggested cell density of 40 – 50 cells per image but the size of the images was not mentioned so comparison of the cell density between different studies is difficult (12).

In the present study, *B. longum* ATCC 15707 culture obtained from early stationary-phase was used for assessing *Bac*Light viability staining because of its relatively stable physiological status (2). It is also desirable to report the proportion of STYO 9 and PI dyes (0.167mM: 1.83mM) for staining approximate 10^8^ cells mL^-1^ of *B. longum* ATCC 15707 in our study. This is because PI can quench SYTO 9 emissions in PI and SYTO 9 stained *Bacillus clausii* (4). SYTO 9 enters the cells of gram positive bacteria (such as *B. longum* ATCC 15707) more easily than gram negative bacteria due to the outer membrane present in the later bacteria (2).

*Bac*Light staining is a useful and efficient method for enumerating *B. longum* ATCC 15707 cells in a pure culture, but after the cells were incorporated into foods, a high background induced by the components or physical fragments from oat and peanut became a challenge when determining green and red fluorescent cells. A real-time viability standard curve of microencapsulated probiotics has been reported using confocal microscopy, but this method might be not precise if encapsulating materials (or food matrices) auto-fluorescence (15).

Sample preparation and analytical methods for assessing viability of probiotics in liquid foods have been reported. Probiotic non-dairy drink samples were filtered through 41 μm and 20 μm filters to remove large solid particles and then centrifuged at 1000 x g for 10 min to concentrate cells, and finally diluted for fluorescence staining (13). Fermented oat drink inoculated with *B. lactis* or a combination of *B. longum* DSM 14579 and *B. longum* DSM 14583 was centrifuged at 800 × g for 7 min to separate a less turbid upper fraction containing cells (9). Although those approaches appeared to achieve satisfying separation results, damage to cells during the separation processes is not known. This might cause underestimation of viability of bacteria that recovered from the stresses present during the separation processes. In the current study, PBDC was applicable to separate *B. longum* ATCC 15707 cells in suspension with food fractions for which differences in buoyant density, thus diminishing fluorescence background and improving accuracy of QFM. This method was validated and confirmed a viability loss of less than log 0.2 cells mL^-1^ after PBDC. It suggests that application of PBDC to separate a probiotic bacterium from solid non-dairy foods such as oat, peanut and raisin is feasible. It is worth considering the limitations of PBDC method where separation process is complex and time-consuming, and this method must be re-examined if other bacteria and food matrices are used, e.g. incorporation into rather than onto food matrices.

An alternative fixation method has been reported by other, bioadhesive slides (Excell AdhesionTM, Fisher Scientific) which showed good cell trapping, but compounds present on the slides caused mortality or affected permeability of the cell membrane, thus inducing mortality of some pure cultures and river water microbes (5). Recently, a low-melting point agarose gel supplemented with *Bac*Light stains was used to rehydrate and determine viability of dried cell particles deposited on Anopore chips (19). However, Anopore chips are expensive for routine determination of live or dead cells in a product. We attempted to use 0.22 μm PC black membrane as an alternative to Anopore chips, but the numbers and ratio of green and red fluorescent cells after transferring from the PC membrane to water agar (3%) were not consistent, thus affecting viability assessment.

Regarding the difference in viability of *B. longum* ATCC 15707 determined using QFM or plate counting, a lower viability was measured by plate counting than the microscopic method in our study. Another study reported 10-fold less by plate counting compared to microscopic observation (16). In another study of *B. lactis* incorporated into butter, the number of cells determined by plate counting was log 2 units lower than the microscopic method after production and storage for one week at 4 °C (18). The difference in numbers of *B. lactis* between the two methods was log 0.7 units after 1 month of storage at 4 °C, but a great difference of log 6.8 units was found in the numbers of *B. longum* inoculated in the same oat drink products (9). Our study shows that plate counting may not provide precise information on the cellular status of *B. longum* ATCC 15707 on non-dairy foods as this bacterium may enter a dormant state during production and storage, and hence the cell numbers using plate counting were underestimated. Therefore, applying two or more different enumeration methods to determine numbers of probiotic bacteria in food products can provide better understanding of bacterial cell status, thus giving an informative result of bacterial viability, especially at elevated temperature of 50 °C when used to investigate the effect of heat stress on cell membrane of *B. longum* ATCC 15707 coated on different non-dairy-foods. This temperature is regarded as having reversible impacts by melting membrane lipids of *L. bulgaricus* cells (23). The viability results of *B. longum* ATCC 15707 coated on different non-dairy foods and dried at 20 °C and 50 °C support that QFM can mutually supports plate counting. When *B. longum* ATCC 15707 coated foods were treated at a mild temperature for 24 h or at a harsh temperature for 2 h, QFM showed its strength in providing information about the cell membrane status, while plate counting reflected the overall impact of drying. This is in accordance with the statement that cultivation method shows bacteria are either detected as cultivable or not, interpreted as alive or dead cells (8). However, this result cannot help to understand the process of how bacteria cell response to drying conditions.

## MATERIALS AND METHODS

### Preparation of bacterial culture and cell suspension

A fresh culture of *B. longum* ATCC 15707 was prepared by growing in MRS broth supplemented with 0.05% (w/v) of *L*-cysteine (MRSc) at 37 °C for 20 h under anaerobic conditions using CO_2_ generating sachets (2.5L, OxoidTM, ThermoFisher, USA). Optical density at 600 nm (OD_600_) was measured to ensure the cell density reached about 1.0 – 2.0, which represents nominal cell populations of 10^8^ ∼ ^9^ cells mL^-1^.

One milliliter of a fresh culture in MRSc broth was transferred into a 1.5 mL Eppendorf tube and centrifuged at 10, 000 × g for 5 min using an Eppendorf^®^ Minispin^®^ microcentrifuge (Sigma-Aldrich, USA). Cell pellets were suspended in NaCl solution (0.15 M) and washed twice at 10, 000 × g for 5 min. After that, cell pellets were re-suspended in 1 mL of NaCl solution (0.15 M) with the final cell density of approximate10^8^ cells mL^-1^.

### Culture coated non-dairy food samples

A fresh culture was added onto oat, peanut and raisin to make a 5% (v/w) loading and then air dried in a biosafety cabinet for 2 h. After that, triplicate culture coated samples were dispensed equally into two sterile petri dishes (90 × 15 mm). One was placed in an incubator at 50 °C and the other was stored at 20 °C for 24 h. The viability of *B. longum* ATCC 15707 on these foods was determined by both QFM and plate counting at times 0, 2 and 24 h.

### Plate counting enumeration of *B. longum* ATCC 15707

Triplicate samples were weighed and placed in a 50-mL tube, sterile MRSc broth was added to make a 10^-1^ dilution. Samples soaked in MRSc broth for 1 h, followed by vortex mixing for 1 min, and a serial dilution for each sample was then prepared. Aliquots of 0.1 mL were plated on MRSc agar plates and incubated 48 h under anaerobic conditions. Colonies on each agar plate were counted and calculated for mean viable cell numbers in each sample.

### Sample preparation using Percoll buoyant gradient density centrifugation (PBDC)

Different Percoll solutions were prepared by diluting stock isotonic Percoll (Percoll:1.5 M NaCl = 9:1) with 0.15 M NaCl as described (6). Percoll density gradients were prepared with a peristaltic pump (LKB pump P-1, Pharmacia), flow rate of < 1 mL min^-1^ by carefully layering 1 mL of each Percoll solution from high density to low density in a 15-mL conical tube. Culture coated food homogenates were loaded on the top layer of seven layers of buoyant gradients for determining buoyant density for *B. longum* ATCC 15707 and fractions of oat, peanut and raisin.

To investigate the effects of PBDC and storage at 4 °C on viability of *B. longum* ATCC 15707, five replicates of a cell suspension of 1 × 108 cell mL^-1^ were prepared in by adjusting OD_600_ to 0.2. A volume of 0.3 mL of each replicate was loaded on the top of 0.6 mL of Percoll gradient solutions of 1.052 g mL^-1^ and 1.121 g mL^-1^ in a 1.5 mL Eppendorf tube, and centrifuged at 14,500 × g for 5 min at room temperature. Following centrifugation 0.5 mL was carefully taken from the top proportion and diluted 10 times using NaCl solution (0.15 M), and the viability of *B. longum* ATCC 15707 was determined by QFM and plate counting. Additional samples were kept overnight in a fridge of 4 °C followed by *B. longum* ATCC 15707 viability determination using QFM and plate counting.

### Quantitative fluorescence microscopy (QFM)

All microscopy work was performed using an Epi 50i fluorescence microscopy (Nikon, Austria) and LIVE/DEAD *Bac*Light viability staining to ensure that green fluorescent cells (SYTO^®^ 9 stained cells) and red fluorescent cells (PI stained cells) were distinguishable. Cy3-4040C (Semrock) fluorescence was excited with the 531 nm line from a Mercury lamp and collected with a triple band pass dichroic mirror (Semrock # FF562-Di03) and a FF01-593/40 emission filter (Semrock). GFP-3035C (Semrock) fluorescence was excited with the 472 nm line from a Mercury lamp and collected with a triple band pass dichroic mirror (Semrock # FF495-Di03) and a FF01-520/35 emission filter (Semrock). Images were acquired with a Nikon DS-Ri2 cooled-CCD camera controlled with NIS-Element BR 4.5.

An aliquot of 50 µL of each sample was loaded on a well of 6-well cell culture plate (flat bottom, Sigma-Aldrich, USA) with filled with water agar (3%) and covered with coverslips (No. 1 1/2) and mounted in a stage. All images were acquired with a Live-fast mode (640 × 480 pixel) on the microscope equipped with Plan Fluo 100 × NA 1.3 objective lens (calibrated before use) giving a final pixel resolution of 0.19 μm, thus the area of the microscopy field of view was 5.8 × 10^4^ μm^2^. Depth of coverslip was 1.7 × 10^1^ μm according to the manufacture instruction, a volume of one microscopy field of view was 9.86 × 10^5^ μm^3^ representing 9.86 × 10^-7^ mL as 1 μm^3^ is equal to 10^-12^ mL. For a direct enumeration of bacteria suspended in one mL of a solution, a microscopic factor of 9.86 × 10^-7^ was used.

Image analysis was performed on QFM images using Image J software (version 1.42q, National Institutes of Health, USA). Images of stained cells collected from single channels (green or red fluorescent) were grouped into green or red cell stacks, and then using MaxEntropy thresholding to distinguish cells.

### Optimal conditions of QFM

washed cell pellets of *B. longum* ATCC 15707 were suspended in 0.15 M NaCl solution to yield a range of cell densities between 10^6^ and 10^9^ cells mL^-1^ as measured by OD_600_ of cell suspensions. Fifty microliters of each cell suspension were stained with 2 μL of 10^-1^ diluted component B in a 1.5 mL Eppendorf tube and mixed properly by pipetting up and down for 30 times. Fifty microliters of *Bac*Light stained cell suspensions were transferred onto a well of a 6-well plate previously filled with water agar (3%), and then spread evenly using an inoculation loop with folded top. Stained cells on the agar plate were placed in the dark immediately and kept at room temperature for 20 min. An optimum range of cell density and proportions of *Bac*Light STYO 9 and PI dyes were determined to give a high fluorescence density of green or red fluorescent cells and a low fluorescence background.

### Accuracy and repeatability of QFM

To obtain 100% dead cells, 1 mL of fresh MRSc culture of *B. longum* ATCC 15707 was heated using a regular laboratory heat block at 85 °C for 15 min (13). The rest of the fresh culture was stored at 4 °C for use as 100% live cells. After 15 min heating, the heated culture was cooled down to room temperature. Later, 100% of live cells and 100% dead cells in MRSc broth were washed and resuspended in 0.15 M NaCl solution to make a concentration of 1 × 108 cell mL^-1^ by adjusting its OD_600_ to 0.2. Finally, five different concentrations of live/dead cells (0, 10, 50, 90, 100%) were prepared by mixing different proportions of washed live and dead cell suspensions in 1.5 mL sterile Eppendorf tubes, and calibration curves was plotted by automatic cell counts and manual cell counts in Image J.

To investigate repeatability of QFM a fresh MRSc culture of *B. longum* ATCC 15707 was prepared in triplicate on three consecutive days. The culture each day was used to prepare six replicates, and the cell density of each replicate was adjusted to 1 × 10^8^ cell mL^-1^. Intra-day and inter-day repeatability was evaluated by determining numbers of green and red fluorescent cells of six replicates for each culture.

### Use of QFM and PBDC for evaluating viability of *B. longum* ATCC 15707 on food samples during 24 h drying at 20 and 50 °C

A systematic demonstration of using PBDC to prepare samples was shown in Fig. 5. Each sample were firstly prepared using the same procedure as plate counting enumeration for making a10^-1^ diluted homogenate. Ten milliliter of sample homogenate was transferred into another 15 mL conical tube and then centrifuged at 1, 880 × g for 5 min. Pellets were discarded, and 10 mL of supernatants was placed in a 15 mL conical tube and centrifuged at 4,800 × g for 15 min. Supernatant was discarded and the pellet was suspended in 1 mL of NaCl solution (0.15 M). The sample suspension was then transferred into a 1.5 mL Eppendorf tube and centrifuged at 10, 000 × g for 5 min. The pellets were washed twice with NaCl solution (0.15 M) at 10, 000 × g for 5 min. After that, the pellets were resuspended in 0.1 mL of NaCl solution (0.15M) in a 1.5 mL conical tube and used as a sample for PBDC.

**FIG 5.**
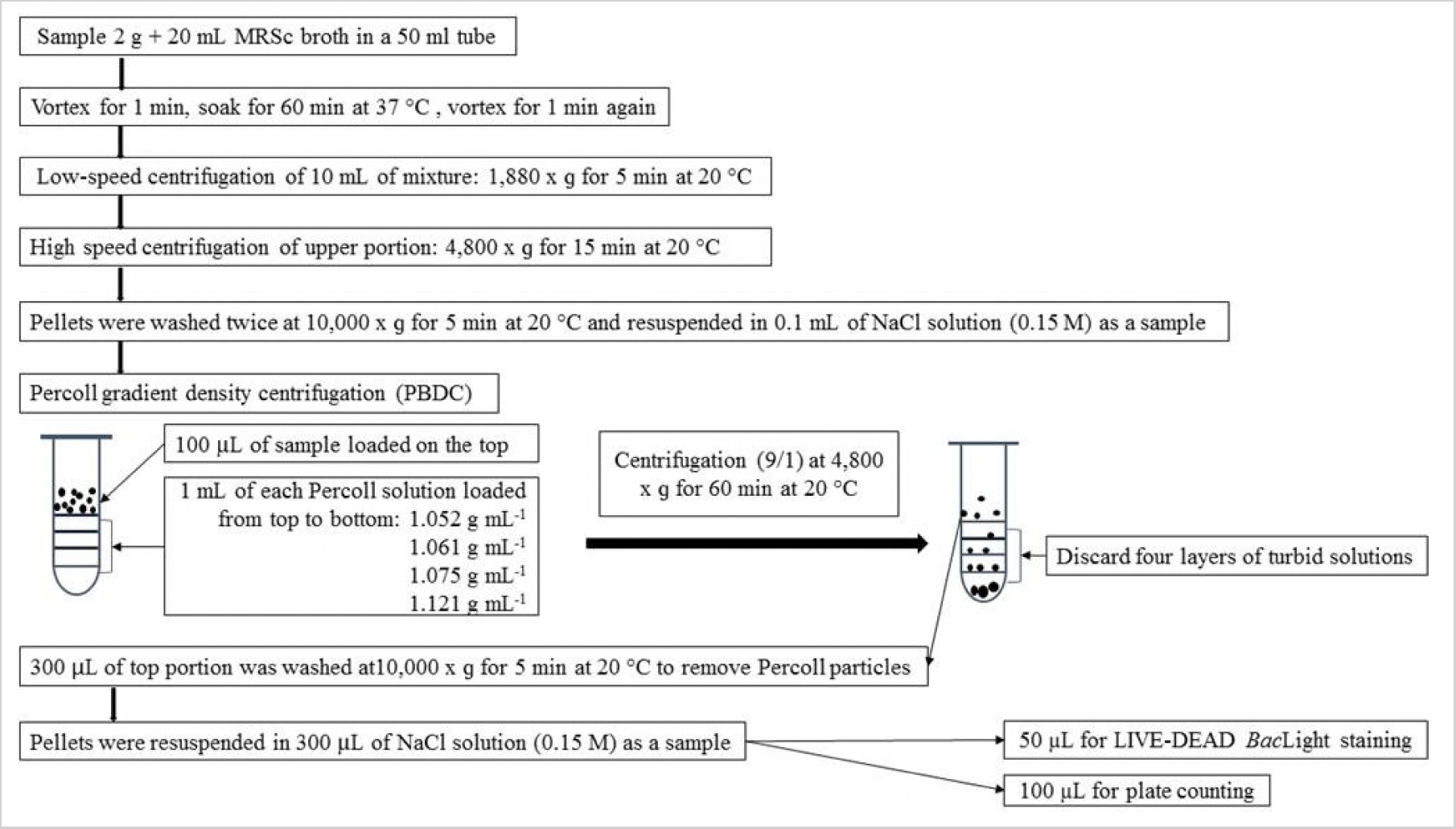
Systematic demonstration of sample preparation by Percoll buoyant gradient density centrifugation (PBDC).

One hundred microliter of the sample solution was placed on top of four x 1 mL layers of Percoll solutions in order (top to bottom: 1.052 g mL^-1^, 1.061 g mL^-1^, 1.075 g mL^-1^ and 1.121 g mL^-1^) in a 15-mL conical tube. The tube was centrifuged at 4,800 × g, Acc/Dcc (9/1) for 60 min. After centrifugation 300 µL of the top portion was carefully transferred to a 1.5 mL Eppendorf tube and diluted with 700 µL of NaCl solution (0.15 M). Percoll particles were removed by centrifuging the sample suspension at 10, 000 × g for 5 min, followed by resuspending the pellets in 0.3 mL of NaCl solution (0.15M) to make a sample for Live/Dead *Bac*Light bacterial viability staining and plate counting. All centrifugation processes were controlled at 20 °C.

